# Structural basis for differential p19 targeting by IL-23 biologics

**DOI:** 10.1101/2023.03.09.531913

**Authors:** Stefano G. Daniele, Sherif A. Eldirany, Minh Ho, Christopher G. Bunick

## Abstract

**Background:** IL-23 is central to the pathogenesis of psoriasis, and is structurally comprised of p19 and p40 subunits. “Targeted” IL-23 inhibitors risankizumab, tildrakizumab, and guselkumab differ mechanistically from ustekinumab because they bind p19, whereas ustekinumab binds p40; however, a knowledge gap exists regarding the structural composition of their epitopes and how these molecular properties relate to their clinical efficacy.

**Objectives:** To characterize and differentiate the structural epitopes of the IL-23 inhibitors risankizumab, guselkumab, tildrakinumab, and ustekinumab, and correlate their molecular characteristics with clinical response in plaque psoriasis therapy.

**Methods:** We utilized epitope data derived from hydrogen-deuterium exchange studies for risankizumab, tildrakizumab, and guselkumab, and crystallographic data for ustekinumab to map drug epitope locations, hydrophobicity, and surface charge onto the IL-23 molecular surface (Protein Data Bank ID Code 3D87) using UCSF Chimera. PDBePISA was used to calculate solvent accessible surface area (SASA). Epitope composition was determined by classifying residues as acidic, basic, polar, or hydrophobic and calculating their contribution to epitope SASA. Linear regression and analysis of variance was performed.

**Results:** All the p19-specific inhibitor epitopes differ in location and size, with risankizumab and guselkumab having large epitope surface areas (SA), and tildrakizumab and ustekinumab having smaller SA. The tildrakizumab epitope was mostly hydrophobic (56%), while guselkumab, risankizumab, and ustekinumab epitopes displayed >50% non-hydrophobic residues. Risankizumab and ustekinumab exhibited acidic surface charges, while tildrakizumab and guselkumab were net neutral. Each inhibitor binds an epitope with a unique size and composition, and with mostly distinct locations except for a 10-residue overlap region that lies outside of the IL-23 receptor epitope. We observed a strong correlation between epitope SA and PASI-90 rates (R^2^ = 0.9969, *p* = 0.0016), as well as between epitope SA and K_D_ (R^2^ = 0.9772, *p* = 0.0115). In contrast, we found that total epitope hydrophobicity, polarity, and charge content do not correlate with clinical efficacy.

**Conclusions:** Structural analysis of IL-23 inhibitor epitopes reveals strong association between epitope SA and early drug efficacy in plaque psoriasis therapy, exemplifying how molecular data can explain clinical observations, inform future innovation, and help clinicians in specific drug selection for patients.

## Introduction

Along with tumor necrosis factor-alpha (TNF-α) and interleukin (IL)-17, IL-23 plays a central role in the pathogenesis of psoriasis (1). Structurally, the IL-23 cytokine is a heterodimer made of the p19 and p40 subunits, which bind IL-23R and IL-23Rβ1 co-receptors, respectively, to activate canonical IL-23 signaling (**Figure 1a**) (2). Importantly, p19 is unique to IL-23, while p40 is shared among other cytokines like IL-12 (3). “Targeted” IL-23 inhibitors risankizumab, tildrakizumab, and guselkumab differ mechanistically from ustekinumab because they bind specifically to p19, whereas ustekinumab binds p40 enabling both anti-IL-23 and anti-IL-12 activity. By binding these distinct subunits, ustekinumab and IL-23-targeted inhibitors exert their therapeutic function by interrupting specific subunit/co-receptor interactions.

**Figure 1.**
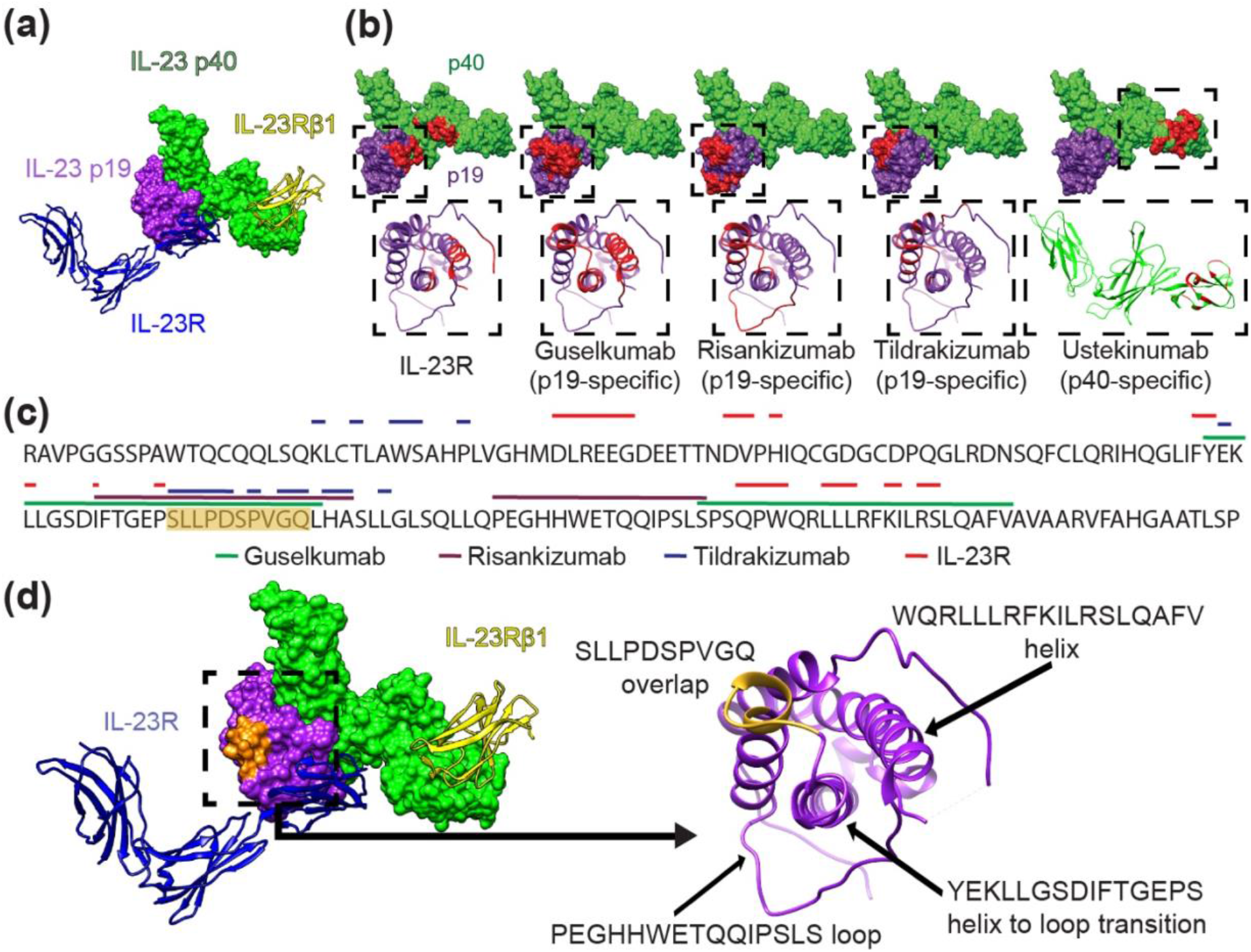
Structural analysis of the IL-23 receptor complex and IL-23 inhibitor binding locations. **(a)** Canonical IL-23 signaling consists of IL-23 p19 (purple) binding to the IL-23R receptor (blue ribbon) and the IL-23 p40 domain (green) binding the IL-23Rβ1 receptor (yellow ribbon). Non-canonical receptor activation may also occur through IL-23R–p19 interaction alone (11). (**b**) IL-23R and IL-23 inhibitor binding sites shown in red on the IL-23 molecular surface (above); IL-23 p19 (purple) and IL-23 p40 (green). Ribbon structures (below) of boxed areas (above) of the p19 or p40 subunits with epitopes colored red. (**c**) The amino acid sequence of the IL-23 p19 subunit is shown with overlying lines corresponding to IL-23R and drug binding sites (ustekinumab is not included because it binds p40). The sequence S95-Q104 containing significant epitope overlap among the IL-23 inhibitors is colored orange, notably in an area that is not a direct IL-23R target. (**d**) IL-23 molecule (left) and ribbon structures of the IL-23R receptor (blue) and of the IL-23Rβ1 receptor (yellow) with the S95-Q104 sequence highlighted from (**c**) in orange. p19 ribbon structure (right) is associated with the boxed area (left) demonstrating the S95-Q104 overlap and other loop and helix secondary structures of p19 that define important epitope components for inhibitors in (**b, c**).

We previously analyzed the binding sites/epitopes of TNF-α, IL-17, and IL-23 inhibitors, identifying molecular differences among the epitopes of drugs within each biologic class and possible links between epitope structure and clinical efficacy (3). Our previous IL-23 inhibitor analysis, however, was limited to ustekinumab because structure data was lacking for other drugs. Hence, structural features of binding epitopes of other IL-23 inhibitors in clinical use have not been closely examined.

Here, we characterize the molecular and structural properties of IL-23 inhibitors by analyzing experimentally-derived epitope data for risankizumab, tildrakizumab, and guselkumab, and hypothesize that structural differences between their epitopes underlie variations in their clinical effects.

## Results

### Epitope location

The epitope of each drug and the binding site of IL-23R were mapped onto the surface of IL-23 (**Figure 1b**). Guselkumab, risankizumab, and tildrakizumab epitopes exist exclusively on p19, while ustekinumab binds exclusively p40 (**Figure 1b**). Risankizumab (2400-Å^2^) and guselkumab (2240-Å^2^) have similar epitope surface areas (SA), while tildrakizumab (1290-Å^2^) and ustekinumab (1390-Å^2^) bind a significantly smaller epitope SA (**Figure 1b**). Importantly, p19-specific inhibitor epitopes differ in location (**Figure 1b**), with guselkumab having the most overlap with the IL-23R binding site: of 24 residues comprising the IL-23R epitope, guselkumab overlaps with 14/24 (58%) residues, risankizumab with 2/24 (8%), and tildrakizumab with 0/24 (0%) (**Figure 1b, c**). One region exists in the IL-23 p19 sequence where all 3 epitopes overlap, specifically 10 residues from S95-Q104 (SLLPDSPVGQ) (**Figure 1c, d**), but this area does not overlap with the IL-23R binding site, suggesting that steric effects—rather than direct competition for the receptor binding site—is sufficient to disrupt receptor binding.

### Epitope chemistry

The chemical composition of the drug epitopes demonstrated a greater hydrophobic character than the IL-23R binding site (32% hydrophobic) (**Figure 2a, b**). Tildrakizumab was the only drug with an epitope having a majority of hydrophobic solvent accessible surface area (SASA) (56%), while guselkumab, risankizumab, and ustekinumab have epitopes with >50% non-hydrophobic residues (polar, acidic, or basic) (**Figure 2b**). Of the p19 epitopes, risankizumab is the only one with a strongly acidic surface charge, while tildrakizumab and guselkumab are net neutral due to relatively even acidic and basic charge distributions (**Figure 2b, c**). On p40, ustekinumab has a strongly acidic epitope (**Figure 2b, c**). For comparison, IL-23R binds a largely neutral surface on p19, but a more acidic region on p40 (**Figure 2c**).

**Figure 2.**
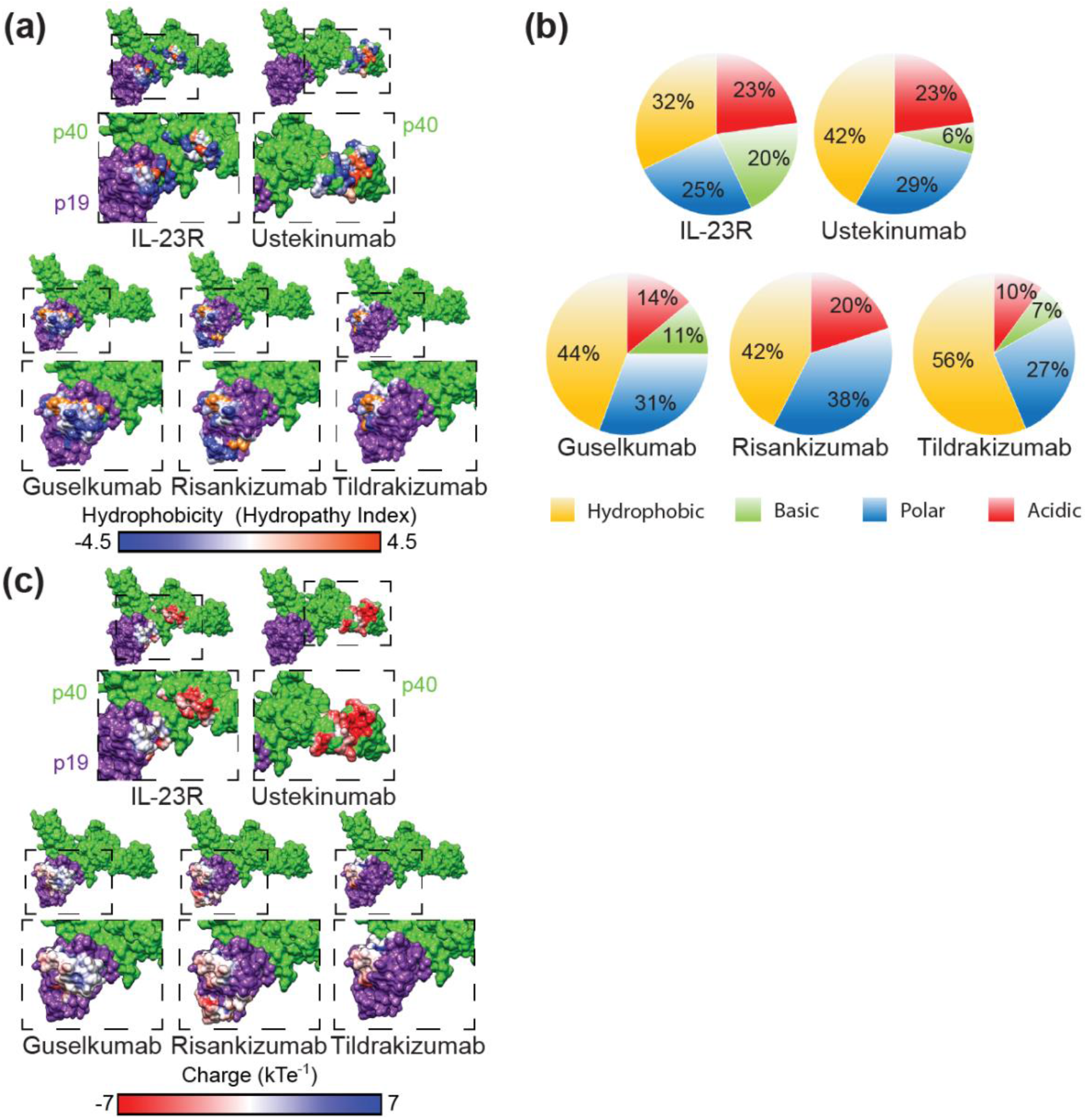
IL-23 inhibitor epitope characteristics. (**a**) Epitope surface hydrophobicity mapped onto IL-23. Polar epitope surfaces are shown in blue, neutral in white, and hydrophobic in orange. Corresponding areas of the dashed boxes (above) are enlarged below. (**b**) Proportion of each total epitope surface accessible surface area (SASA) made up of acidic, basic, polar, or hydrophobic amino acids. (**c**) Epitope surface charges mapped onto IL-23. Acidic epitope surfaces are shown in red, neutral in white, and basic in blue. (**a, c**) IL-23 p19 (purple) and IL-23 p40 (green). Corresponding areas of the dashed boxes (above) are enlarged below.

### Correlation of epitope molecular properties with clinical response

A study comparing IL-23 inhibitor efficacy showed that risankizumab and guselkumab demonstrated high rates of reduction in Psoriasis Area Severity Index-90 (PASI-90), whereas tildrakizumab had a lower rate of achieving PASI-90 (4). We compared these data (4) with the structural properties of each epitope uncovered during our analyses. In doing so, we found a stark correlation between epitope SA of each drug and PASI-90 rates: drugs with higher epitope SA (risankizumab and guselkumab) demonstrated higher short-term (10-16-weeks) PASI-90 rates, while tildrakizumab, which exhibits the smallest epitope SA, demonstrated the lowest rate of PASI-90 over the same timeframe (**Figure 3a**; R^2^ = 0.9969, *p* = 0.0016). We found a similar correlation between epitope SA and short-term PASI-100 scores (R^2^ = 0.9986; data not shown). Notably, ustekinumab fits within this trend, indicating that ustekinumab’s dual IL-12/IL-23 inhibition may not cause it to behave differently from the targeted IL-23 inhibitors; rather, its efficacy may be due to the size of its epitope SA. We next sought to examine whether the drug epitope SA correlated with its known equilibrium dissociation constant (K_D_) (5). Linear regression analysis demonstrated a significant inverse relationship between epitope SA and K_D_ (**Figure 3b**; R^2^ = 0.9772, *p* = 0.0115), indicating that increased epitope SA reflects higher epitope binding affinity. As expected, increased binding affinity also translated to higher PASI-90 rates (**Figure 3c**; R^2^ = 0.9908, *p* = 0.0046). Interestingly, total residue hydrophobicity (R^2^ = 0.3941, *p* = 0.3722), polarity (R^2^ = 0.7369, *p* = 0.1416), and charge (R^2^ = 0.005, *p* = 0.9779) content did not correlate with clinical efficacy as measured by PASI-90 rates (**Figures 3d-f**). However, these correlations are based on short-term clinical responses only, and further studies are needed to assess the validity of these associations for long-term clinical responses.

**Figure 3.**
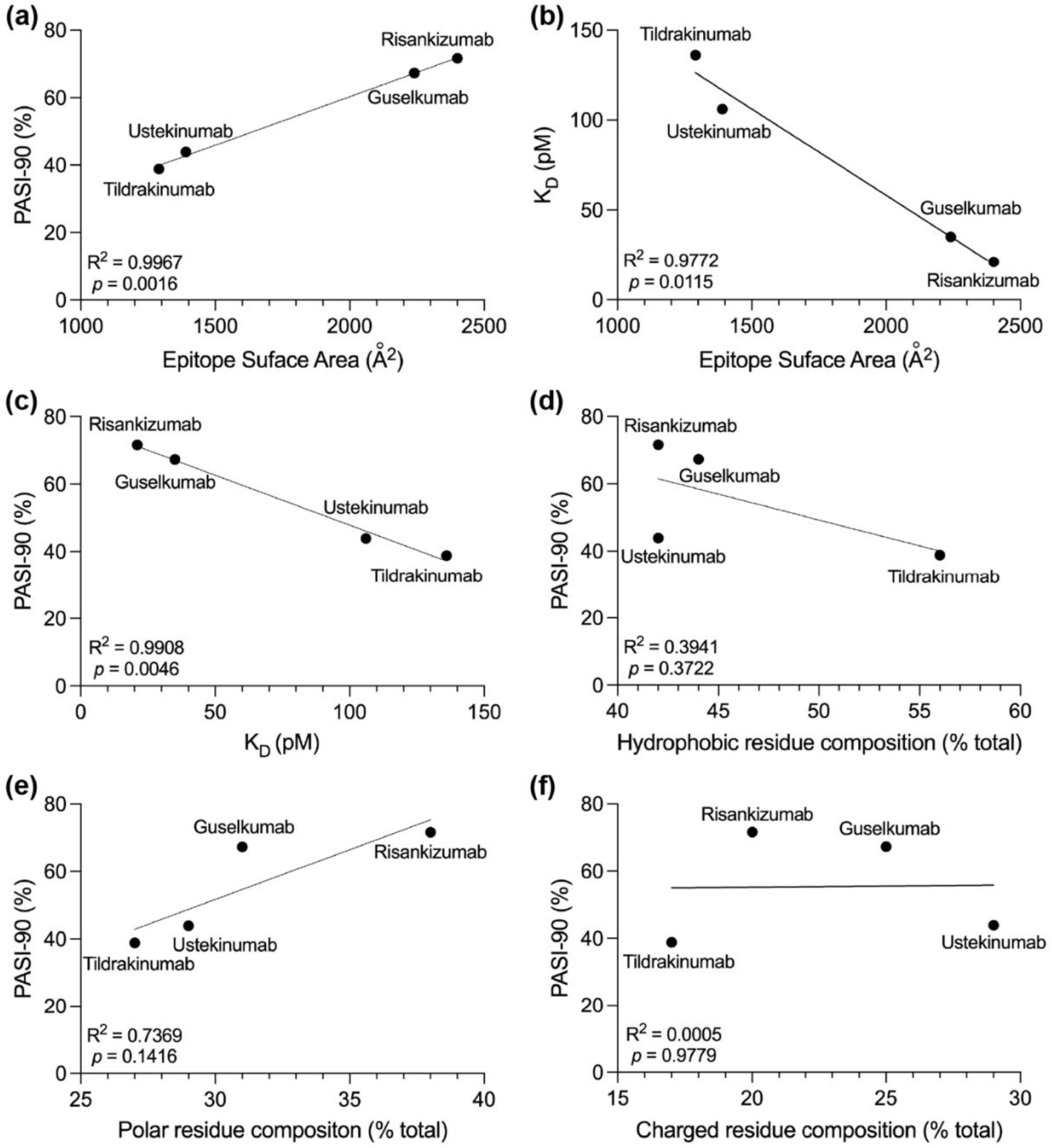
IL-23 inhibitor epitope surface area correlates with clinical efficacy and binding affinity. (**a**) The efficacy of each IL-23 targeting drug, as measured by 10-16-weeks PASI-90 response by patients, is directly correlated with the size of the epitope surface area bound on IL-23. For tildrakizumab: epitope surface area = 1290 Å^2^, PASI-90 = 38.8%; ustekinumab: epitope surface area = 1390 Å^2^, PASI-90 = 43.9%; guselkumab: epitope surface area = 2240 Å^2^, PASI-90 = 67.3%; risankizumab: epitope surface area = 2400 Å^2^, PASI-90 = 71.6%. PASI-90 data derived from (4). (**b**) The K_D_ of single-chain human IL-23 is inversely correlated with the size of the epitope surface area. K_D_ values are derived from (5). (**c**) The K_D_ of single-chain human IL-23 is inversely correlated with PASI-90. (**d**-**f**) Total residue hydrophobicity, polarity, and charge composition do not correlate with PASI-90 clinical response.

## Discussion

Deep examination of available structural data describing IL-23 inhibitor epitopes reveals differences that have real-world implications for clinical practice. Strong correlation between epitope SA and early drug efficacy in plaque psoriasis exemplifies how molecular data help to explain clinical observations and may inform future pharmacologic innovation. Moreover, our data provide clinicians with a molecular rationale for helping them in drug selection among the various IL-23 inhibitors to optimize outcomes for patient care. This information may be especially useful in circumstances when clinicians and patients engage in shared decision-making to trial a different IL-23 inhibitor due to poor clinical response with a prior inhibitor.

Our data are in accordance with previous preclinical studies examining the thermostability, IL-23 binding affinity, as well as the *in vitro* potency and *in vivo* efficacy of risankizumab, tildrakizumab, guselkumab, and ustekinumab (5). In the present study, we deepen the molecular understanding of these IL-23 inhibitors by characterizing and differentiating their structural epitopes. In doing so, we provide the structural rationale underpinning not only pre-clinical findings (5), but also the short-term clinical efficacy of these medications in the treatment of plaque psoriasis (4). Moreover, our data confirms that direct IL-23 inhibitor epitope overlap with the IL-23R epitope is not required for clinical efficacy, since risankizumab and tildrakizumab demonstrate clinically meaningful PASI-90 responses despite low (8% and 0%, respectively) epitope overlap with IL-23R. Indeed, we also find that each IL-23-targeted inhibitor epitope corresponds to a unique location on the IL-23 p19 subunit except for a 10-residue region from S95-Q104 (SLLPDSPVGQ). Importantly, this area likewise does not overlap with the IL-23R binding region, providing further evidence that direct binding interference between inhibitor and IL-23R is not necessary for clinical efficacy.

Our ability to connect structure and function is limited by the lack of detailed drug-cytokine structures for the medications presented herein; the current work relies mostly on biochemical, non-crystallographic data acquired from hydrogen-deuterium exchange experiments. Future studies yielding experimentally-determined structures, from X-ray crystallography or Cryo-electron microscopy, can enhance precision medicine in dermatology by deciphering the molecular and structural mechanisms governing biologic drug activity.

## Methods

### IL-23 epitope analysis

We utilized epitope data primarily derived from hydrogen-deuterium exchange studies for risankizumab (6), tildrakizumab (7), and guselkumab (5), and crystallographic data for ustekinumab (8) to map epitope locations, hydrophobicity, and surface charge onto the surface of the IL-23 crystal structure (Protein Data Bank (PDB) ID Code 3D87) (9) using UCSF Chimera (10). PDBePISA was used to calculate SASA. Total epitope characters were determined by classifying each epitope residue as acidic (D, E), basic (R, K), polar (Q, N, H, S, T, Y, C, G), or hydrophobic (A, I, L, F, V, P, M, W) and calculating each group’s total contribution to epitope SASA.

### Statistical Analysis

All statistical analyses (linear regression analysis) and data plotting were performed using GraphPad 9 (GraphPad Software, Inc.) with significance set at *P* ≤ 0.05. Data utilized for each analysis are described in detail in corresponding figure legends.

## Acknowledgements

We thank Dr. Matt Vesely (Yale University) for critical review. Work supported by NIH/NIGMS Medical Scientist Training Grant T32GM007205 (SGD). Aspects of work were presented at 2022 South Beach Symposium, Miami, FL (CGB).

## Author Contributions

Conceptualization, CGB; analysis, SGD, SAE; writing—original draft preparation, SGD, SAE; writing—review and editing, SGD, CGB; visualization, SAE, SGD, MH, CGB; supervision, CGB. All authors have read and agreed to the published version of the manuscript.

